# Structural investigation of an engineered feruloyl esterase with improved MHET degrading properties

**DOI:** 10.64898/2026.01.05.697840

**Authors:** Panagiota Karampa, Konstantinos Makryniotis, Theofani-Iosifina Sousani, Evangelos Topakas, Vangelis Daskalakis, Maria Dimarogona

**Affiliations:** Laboratory of Structural Biology and Biotechnology, Department of Chemical Engineering, University of Patras, Greece; Industrial Biotechnology & Biocatalysis Group, Biotechnology Laboratory, School of Chemical Engineering, National Technical University of Athens, Greece; Laboratory of Biomolecular Dynamics and Engineering, Department of Chemical Engineering, University of Patras, Greece; Institute of Chemical Engineering Sciences (FORTH/ICE-HT), Foundation for Research and Technology Hellas, Stadiou St, Platani, GR26504 Patras, Greece

**Keywords:** feruloyl esterase, G122S mutation, MHETase, X-ray crystallography, molecular dynamics simulations, docking simulations, MHET

## Abstract

MHETases are enzymes implicated in polyethylene terephthalate (PET) biodegradation. The present study elucidates the structural determinants that result in increased mono(2-hydroxyethyl) terephthalate (MHET) degradation by a feruloyl esterase, which has been engineered to resemble MHETase active site. The crystal structures of the variant in apo and benzoic acid bound state reveal the changes induced by the introduced mutation, specifically the formation of a hydrogen bond and a *trans* to *cis* isomerization of a peptide bond in the vicinity of the catalytic site. Molecular dynamics simulations demonstrate the stabilization of the loop harboring the engineered residue, as well as an expansion of the substrate binding cleft, which would facilitate accommodation of a broader variety of substrates, indicative of a promiscuous biocatalyst.

**Enzyme:** Enzyme Commission Number: EC 3.1.1.73; Uniprot accession code: A0A1D3S5H0_FUSOX

## Introduction

Plastic accumulation in the biosphere has severe impact on the environment and human health. As a result, intensive research efforts have lately focused on the sustainable decomposition and recycling of plastics, employing chemical, mechanical and biological approaches. Regarding the latter, the enzymatic depolymerization of PET is accomplished via the combined use of two enzyme categories: PETases, that cleave the polymeric chains and MHETases, that cleave PETase products, specifically bis(2-hydroxyethyl) terephthalate (BHET) and mono(2-hydroxyethyl) terephthalate (MHET), to terephthalic acid (TPA) and ethylene glycol (EG) [1]. While research on PETases has surged in recent years, studies on the biochemical and structural characterization of MHETases remain quite limited. Structurally, MHETases resemble tannase-like ferulic acid esterases (FAEs, EC 3.1.1.73), which are enzymes that cleave the ester bond between hydroxycinnamic acids and sugars in the plant cell wall [2]. They comprise a hydrolase domain and an α-helical lid domain, stabilized by a calcium ion. Structural studies of MHETase from *Ideonella sakaiensis*, namely *Is*MHETase, in complex with PET oligomers have allowed the identification of amino acids that mediate MHET binding in the active site and the description of MHET degradation mechanism [3], [4], [5], [6].

*Fo*FaeC is a tannase-like FAE from *Fusarium oxysporum* that has been biochemically and structurally characterized [7], [8]. It is a serine-type hydrolase, comprised of a catalytic and a lid domain with a calcium binding site, similarly to *Is*MHETase. *Fo*FaeC catalytic triad consists of Ser201, His452 and Asp412, while the amide groups of Gly123 and Thr202 form the oxyanion hole [7]. Due to its structural homology to MHETase, its potential exploitation as plastic degrading enzyme was investigated. Thus, an *Fo*FaeC variant, namely *Fo*FaeC_G122S, designed in order to mimic MHETase active site, was shown to exhibit improved MHET and BHET degrading activity, compared to the unmodified enzyme, without compromising its thermostability [9]. When tested on additional synthetic model substrates it was found that the variant was capable of hosting bulkier PET analogues, such as M(HET)_2_, a soluble methylated PET dimer, in contrast to wild-type (WT). The enzyme also displayed improved catalytic properties against methyl esters of hydroxycinnamic acids, specifically methyl caffeate (MCA) and methyl *p*-coumarate (M*p*CA), while exhibiting reduced selectivity among them, in contrast to WT which preferentially cleaves methyl ferulate (MFA) [9]. Thus, it seems that mutation G122S results in a more promiscuous enzyme, without affecting its efficiency as biocatalyst. Here, the crystal structure of *Fo*FaeC_G122S variant is presented, in apo and benzoic acid (BA) bound state. Analysis of the determined structures highlights the introduced intramolecular interactions and structural alterations that append these improved properties to the engineered enzyme. Further docking and molecular dynamics simulations, of apo and MHET bound enzymes, corroborate the experimental findings, contributing to a better understanding of the structure-function relationships of plastic degrading enzymes, with potential for biotechnological applications.

## Materials and Methods

### Cloning, expression, and purification

Recombinant *Fo*FaeC_G122S was expressed in *Komagataella pastoris* and purified using immobilized metal-ion affinity chromatography with Co^2+^ (Talon; Clontech Laboratories Inc., CA, USA) as described previously [9]. The chromatography column was equilibrated with 20 mM Tris-HCl buffer containing 300 mM NaCl (pH 8.0). After loading the sample, it was sequentially washed with 10 mL buffer, 7 mL of 5 mM imidazole and 7 mL of 10 mM imidazole in 20 mM Tris-HCl containing 300 mM NaCl (pH 8.0). The protein was eluted with 3 consecutive stages of 6 mL of 100 mM imidazole in the same buffer. A final polishing step was performed by size-exclusion chromatography (SEC) on a 16/60 Sephacryl column. The purity of *Fo*FaeC_G122S was checked by sodium dodecyl sulfate-polyacrylamide gel electrophoresis (SDS-PAGE) with a single band corresponding to a molecular mass of 70 kDa.

### Crystallization and structure determination

Purified *Fo*FaeC_G122S was concentrated to 16 mg·mL^−1^ in 20 mM Tris-HCl pH 8.0 and submitted to crystallization trials using sitting drop vapor diffusion technique. Crystallization experiments were conducted with the aid of an OryxNano crystallization robot (Douglas Instruments Ltd, UK). Diffracting plate-like crystals were grown by mixing equal volumes (0.2 μL) of protein and reservoir solution after 2 days in the presence of 0.1 M Tris-HCl pH 8.5 and 35% (v/v) PEG400 at 20 °C (Fig. S1). For co-crystallization, *Fo*FaeC_G122S and MHET were mixed at final concentrations of 15.2 mg·mL^−1^ and 10 mM, respectively, and incubated on ice for 40 min. Afterwards, crystallization was performed as described above, mixing 0.4 μL protein-MHET solution and 0.2 μL reservoir solution. Crystals appeared within 2 days in the presence of 0.1 M Tris-HCl pH 8.5, 37% (v/v) PEG400.

For X-ray data collection, crystals were mounted on litholoops and flash cooled in liquid N_2_. X-ray diffraction data were collected at beamline P13, operated by EMBL Hamburg at the PETRA III storage ring (DESY, Hamburg, Germany) [10]. *Fo*FaeC_G122S and MHET-*Fo*FaeC_G122S crystals diffracted to 1.90 Å and 1.71 Å, respectively. Both crystals were grown in space group *P*2_1_ with 2 molecules in the asymmetric unit. The resulting datasets were processed using *XDS* [11] and the data integration and scaling was performed with *AIMLESS* [12] at the *CCP4i2* program suite [13]. Both crystal structures were determined by molecular replacement with *PHASER* [14] using the apo structure of *Fo*FaeC as starting model (PDB code: 6FAT). Iterative rounds of manual model building and refinement of the structure were also performed using *COOT* [15] and *REFMAC5* [16], respectively. The atomic coordinates and structure factors for crystal structures of *Fo*FaeC_G122S and *Fo*FaeC_G122S in complex with BA have been deposited in the Protein Data Bank (PDB; https://www.rcsb.org/) under accession codes 9I50 and 9HUN, respectively.

### Molecular dynamics (MD) simulations and analysis

The initial coordinates were taken from the *Fo*FaeC crystal structure (PDB code: 6FAT, chain: B) for the wild-type (WT) and *Fo*FaeC_G122S (PDB code: 9I50, chain: B) for the mutant (MT). Glycosylations and other ligands were removed prior to the preparation of the systems, otherwise all resolved water molecules were retained in the structures. The protonation states of Glu and Asp residues were predicted by *PROPKA* method [17], [18] in the *PDB2QR* server at pH 6.0 (https://server.poissonboltzmann.org/) and, also, validated by visual inspection to preserve the hydrogen bonding network within the resolved proteins. In detail, all His residues were protonated only at the N_ε_ sites, except His73, His174, His192, His252 and His452 that were protonated only at the N_δ_ sites. Asp367, Glu210 and Glu346 were kept always protonated, whereas all other acidic residues were left deprotonated. The model structures were embedded in a 12×12x12 nm^3^ simulation box-cell and hydrated by around 55,000 TIP3P water molecules [19]. Each system was neutralized by adding 5 chloride anions (Cl^−^) (∼173k atoms in total/system). To be consistent with the crystallographic conditions, no additional salt ions were added. The Charmm36 Force Field [20] was employed for the description of the polypeptide chains and ions.

Based on published protocols, all models were relaxed and equilibrated with gradual removal of constraints on the protein backbone-heavy atoms [21]. Briefly, a series of constant volume (nVT), and constant pressure (nPT) ensemble runs increased the temperature from 100 K to 308 K over 50 ns prior to production runs. Classical MD simulations were run for 500 ns in 4 independent trajectories per system (WT and MT) with variant initial coordinates-velocities derived from the last step of equilibration (1 ns apart). Newton’s equations of motion were integrated with a time step of 2.0 fs. The leapfrog integrator in *GROMACS 2022.5* was employed [22]. The production runs have been performed in the constant pressure nPT ensemble with isotropic coupling (compressibility at 4.5×10^−5^). Moreover, the Van der Waals interactions were smoothly truncated between 1.0-1.2 nm with the Verlet cut-off scheme (force-switch algorithm and tolerance at 10^−3^). Short-range electrostatic interactions were truncated at 1.2 nm and long-range contributions were computed within the Particle-Mesh-Ewald (PME) approximation with tolerance at 10^−5^ and Fourier spacing at 0.15 [23]. All hydrogen-heavy atom bond lengths were constrained after equilibration employing the LINCS algorithm [24]. For the production, the v-rescale thermostat is employed [25] (temperature coupling constant = 0.5) and the C-rescale barostat [26] (1 atm; pressure coupling constant = 2.0). The first 100 ns of each trajectory were considered as further equilibration time and were disregarded from the analysis. In total, the production sampling time of these 2 systems with 4 trajectories/system running for 0.5 μs each is 4 μs.

For the analysis, Principal Component Analysis (PCA) and k-means clustering were performed by in-house scripts based on the sklearn python library. The Root-Mean-Square Fluctuation (RMSF) and B-factors were calculated for the protein CA atoms and weight-averaged based on the eigenvalues of the three main PCA components per trajectory that represent 35-40% of the total motion sampled per replicate trajectory. Four clusters were identified per system (WT and MT) by the k-means algorithm and the representative structure from each cluster was extracted for docking simulations.

### Molecular Docking of MHET

Docking analysis was performed using *YASARA 21.6.17* software [27]. The key conformations for *Fo*FaeC and *Fo*FaeC_G122S obtained from the MD analysis (k-means clustering) were used as initial coordinates for docking. Water molecules were removed, and energy minimization of all structures was held in *YASARA,* without compromising the identity of the predicted key conformations. MHET was built as follows: a non-hydrolysable form of MHET, named MHETA, captured at the catalytic center of a MHETase was downloaded from PDBe (PDB code: 6QGA) [5] and the nitrogen atom was substituted by oxygen to build the ester bond using *YASARA*. The ligand was placed in the active site of each structure by aligning with MHETase using *MUSTANG* [28] and the resulting structures were submitted to perform local docking in *YASARA*. Structures obtained from docking analysis were ordered based on their binding free energy and predicted dissociation constant (*K*_d_). The docked structure with the most favorable energy and the lowest *K*_d_ of each enzyme was selected for further MD simulations, with the constraint that the distance between the hydroxyl group of the catalytic serine (Ser201) and the carbon atom of the carbonyl group connected to the ethylene glycol moiety of MHET did not exceed 4 Å. *CHARMM-GUI* [29], [30] was used to construct the compatible parameter files of MHET for Charmm36 Force Field. Classical MD simulations were run for 500 ns in 4 independent trajectories per complex system (WT and MT), as in the apo structures.

### Thermostability

Thermodynamic stability of both *Fo*FaeC and *Fo*FaeC_G122S enzymes was checked through essays at 308, 328 and 348 K. Initial coordinates of each system were taken from the last frame (t=500 ns) of the apo MD simulations at 308 K. The structures were heated to the goal temperature using a 10 ns-simulation. Then, the classical MD simulations for each system were run for 100 ns saving the system snapshots every 10 ps.

For the analysis, the dimensionless parameter (Λ) was used, describing the dependance of the squared generalized order parameters (S^2^) for the backbone N-H bond vectors on the temperature [31], [32]. The first residue (Asp36) and all prolines were excluded from the calculation, as Asp36 is charged due to the appearance of three hydrogen atoms connected to the N and prolines lack the presence of hydrogen atom to the nitrogen. The Λ values were determined providing thus information about the temperature dependent fluctuations of N-H bond vectors. Lower Λ values indicate a more thermostable protein molecule.

## Results & Discussion

### Crystal structure of FoFaeC_G122S

The crystal structure of *Fo*FaeC_G122S was determined to a resolution of 1.90 Å, in apo form (PDB code: 9I50), and to 1.71 Å, in complex with BA (PDB code: 9HUN). Data collection and refinement statistics are shown in Tables S1 and S2. Similarly to the unmodified enzyme, *Fo*FaeC_G122S crystallized in space group *P*2_1_ with 2 molecules in the asymmetric unit (a.u.). It is composed of an α/β hydrolase domain, and an α-helical lid domain composed of residues 229-389 (Fig. 1A). The post-translational modifications observed in the WT enzyme are also present in the modified one. Specifically, N-linked glycans at residues 66, 101, 111, 151 and 362, as well as the six disulfide bonds between cysteines 41-88, 76-127, 200-453, 269-287, 296-304, and 516-538 are identified. The Cys200-Cys453 disulfide bond is partially broken, likely due to the radiation damage. In its reduced state, Cys200 forms hydrogen bonds with three water molecules, one of which also interacts with the hydroxyl group of the engineered Ser122 (Fig. 1B). The lid domain is stabilized by a calcium ion that is coordinated by seven oxygen atoms in each monomer, as in the WT.

**Figure 1.**
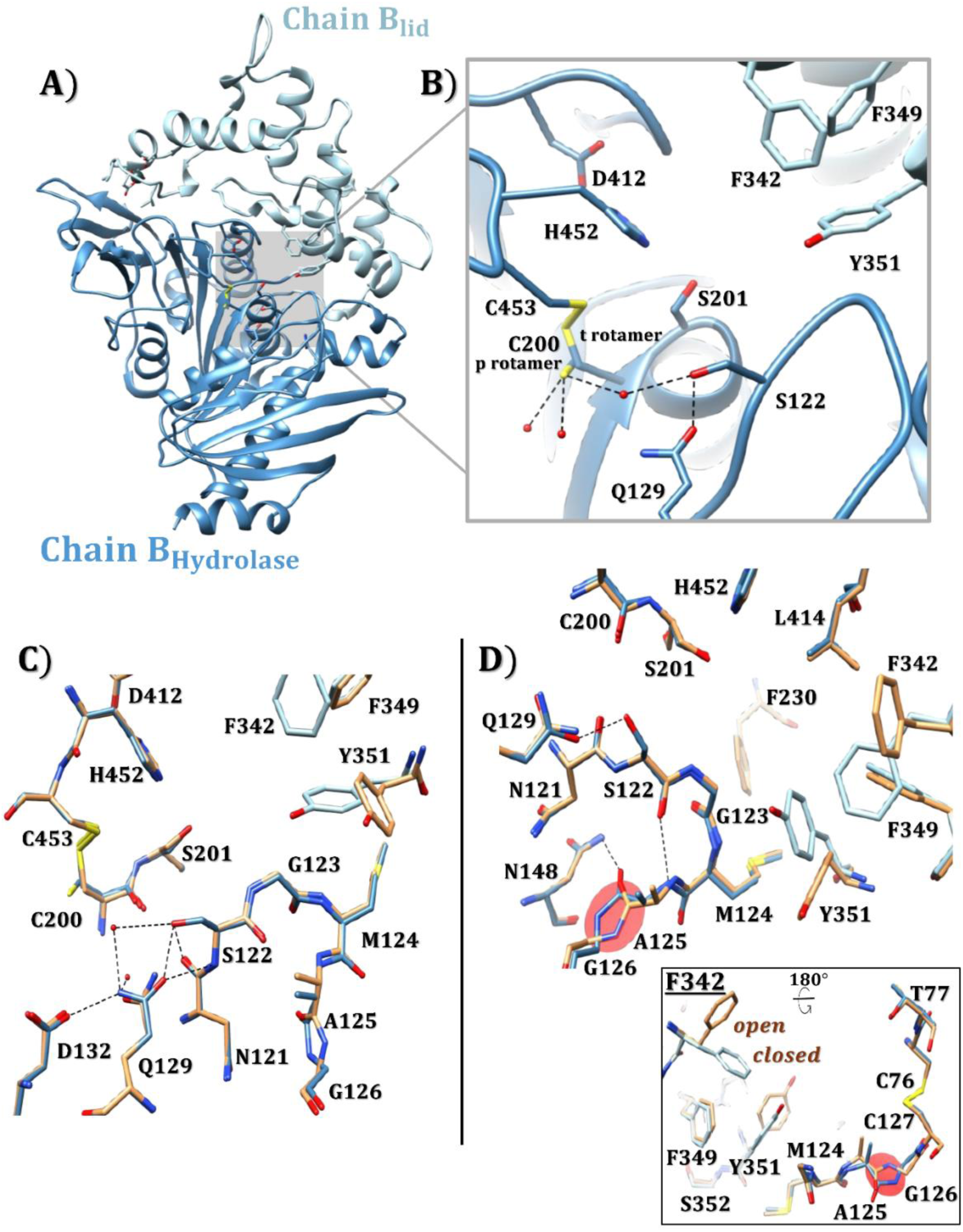
Crystal structure of FoFaeC_G122S. **(A)** Cartoon representation of FoFaeC_G122S monomer (molecule B). Lid domain (residues 229 – 389) is represented in light blue, while the hydrolase domain is in steel blue. Calcium ion is shown in dark gray sphere. **(B)** The two rotamers of Cys200 that participate in disulfide bond formation are shown. The reduced form of Cys200 forms hydrogen bonds (depicted in dashed black lines) with three water molecules (red spheres). Amino acids are shown as sticks, while oxygen and nitrogen atoms are shown in red and blue, respectively. **(C)** Stick representation of FoFaeC_G122S superimposed on the wild-type FoFaeC (PDB code: 6FAT, molecule B). Close-up view of the mutation region of FoFaeC_G122S, with hydrogen bonds formed between the engineered Ser122 and Gln129, as well as between Gln129 and Asp132. **(D)** Stick representation of the cis/trans conformation between Ala125 – Gly126, highlighting the closed/open conformation of Phe342. The hydrogen bonds shown in dashed black lines correspond to the FoFaeC_G122S structure.

Superposition of chain B of WT (PDB code: 6FAT) and *Fo*FaeC_G122S structures over all atoms indicates an RMSD of 0.248 Å. In agreement with that, visual inspection of the superimposed structures does not indicate significant structural alterations. The most evident change induced by the engineered amino acid is the formation of three hydrogen bonds. Two of these involve interactions between Ser122 and Gln129, in which the side chain hydroxyl group and the backbone amide of engineered Ser122 form hydrogen bonds with the side chain carbonyl group of Gln129, inducing a shift of its C_d_ atom by 0.3 Å towards Ser 122 (Fig. 1C). The third hydrogen bond is formed between the NE2 atom of Gln129 and the carboxyl group of Asp132. In contrast, in the unmodified enzyme, Gln129 interacts only with 2 neighboring water molecules, that are also conserved in *Fo*FaeC_G122S. In addition to that, the peptide bond between Ala125 and Gly126 adopts a twisted *trans* conformation in molecule A and a *cis* conformation in molecule B of *Fo*FaeC_G122S, in contrast to the wild-type enzyme, where this bond adopts a *trans* conformation in both monomers in the asymmetric unit. This *cis* conformation is accompanied with a “closed” conformation of Phe342, pointing towards the putative ligand binding site (Fig. 1D). This alteration results in the formation of a hydrogen bond between the carbonyl oxygen of Ala125 and atom ND2 of Asn148.

### Crystal structure of FoFaeC_G122S in complex with benzoic acid

The *Fo*FaeC_G122S-benzoic acid (BA) complex structure reveals a set of noncovalent interactions mediating the binding of the ligand. Specifically, the benzoic carboxyl group forms hydrogen bonds with the hydroxyl group of one of the two rotameric configurations of the catalytic Ser, the side chain hydroxyl and the backbone amide of Thr202, and the backbone amide of Gly123 (Fig. 2B). The two latter amino acids form the oxyanion hole said to stabilize the tetrahedral intermediate [4]. Regarding the side chain of the catalytic Ser, while in the apo form of *Fo*FaeC_G122S it only appears in the t rotameric form, in the BA-bound structure, it is in both t and p rotameric forms [33], which results in its Oγ atom being 2.40 Å and 3.49 Å from the carboxyl oxygen of BA, respectively (Fig. 2B, 2C). The p rotamer is stabilized through a hydrogen bond with the hydroxyl group of engineered Ser122, and is not observed in the *p*-coumaric bound structure of *Fo*FaeC WT, where an m rotameric form appears (Fig. 2C). The aromatic ring of BA is stabilized via T-shaped π-π interactions with Tyr351, in a hydrophobic environment created by Phe230, Phe349, Phe342, Leu414 and Ile415 (Fig 2B). The *cis* conformation of the peptide bond between Ala125 and Gly126 is observed in both molecules of the a.u., similarly to monomer B of “apo” *Fo*FaeC_G122S and both monomers of *Fo*FaeC in complex with *p*-coumaric acid (*p*CA) (PDB code: 8BHH) [8]. In addition, in both monomers, Phe342 adopts an inward conformation. This phenylalanine placement, which appears in tandem with the *cis* Ala125-Gly126 bond, is only observed in substrate-bound structures of the enzyme, as well as molecule B of apo *Fo*FaeC_G122S, as described above. It should be mentioned that unmodeled electron density at the active site of monomer B indicates the occurrence of an unidentified ligand (Fig. S2), probably originating from the crystallization solution, other than water, that could influence the configuration of amino acids forming the binding cleft. Regarding Tyr351, which has been previously shown to change conformation upon substrate binding, and whose mobility upon substrate binding as Phe415 in *Is*MHETase has also been previously demonstrated [5], it does not seem to change conformation between the apo and BA bound state. Still, electron density indicates increased mobility of this amino acid (Fig. S3), further confirmed by its relatively increased B-factors, also observed in the neighboring Pro350 and Phe349. Compared to the *p*CA-bound *Fo*FaeC WT structure, the aromatic rings of the two ligands occupy nearly identical positions, with their C1 carbons overlapping. However, the C4 carbon is rotated by approximately 33° around an axis perpendicular to the ring plane and passes through C1 of the benzene ring (Fig. 2C).

**Figure 2.**
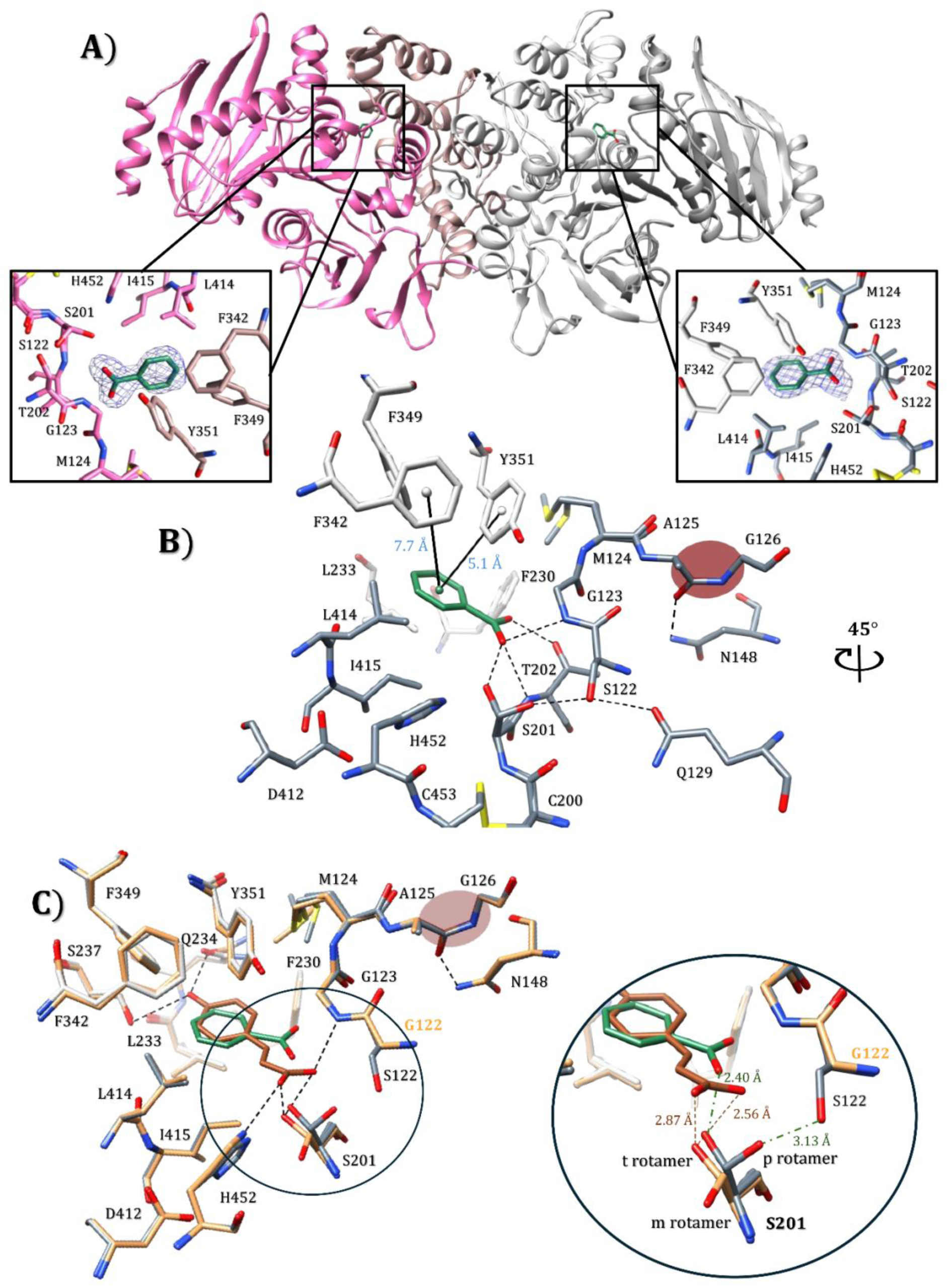
Crystal structure of FoFaeC_G122S in complex with benzoic acid. (A) Cartoon representation of both monomers in the a.u., highlighting the substrate binding site of FoFaeC_G122S with benzoic acid modeled in both molecules (chain A in magenta/sandy brown and chain B in dark/light gray). Electron density maps are contoured at 1σ (blue mesh). Benzoic acid molecules are shown as dark green sticks. **(B)** Close-up view of the substrate binding site. The catalytic triad (Ser201-His452-Asp412) and the hydrophobic environment (Phe230, Leu233, Phe342, Phe349, Tyr351, Leu414, Ile415) are shown as sticks. The hydrogen bonds are shown in dashed black lines, while the distances between the centroids of Phe342/Tyr351 and BA are shown in continued black lines. **(C)** Superposition of FoFaeC_G122S-BA and wild-type FoFaeC-pCA with a close-up view of the binding pocket.

In conclusion, considering also our previous results on *Fo*FaeC apo and *p*CA bound structures, even though Tyr351 displays considerable mobility, the most evident structural rearrangement upon substrate binding includes Phe342 which seems to be implicated to the substrate placement into the active site (Fig. 2B) and the adoption of *cis* conformation in the peptide bond connecting residues 125 and126. The distance between Phe342-BA centroids is 7.7 Å (Fig. 2B, 4B), which could not imply a direct π-π interaction [34].

### Comparison with BA bound IsMHETase structure

Currently, there are two crystal structures of *I. sakaiensis* MHETase in complex with BA (PDB codes: 6QGB, 6QZ3), both with BA orientation almost opposite to *Fo*FaeC_G122S-BA structure (Fig. 3A). This position arises from the ionic interaction between the carboxylate group of BA and the positively charged Arg411 in *Is*MHETase, which is not conserved in *Fo*FaeC. The importance of Arg411 for TPA recognition via its positive charge has been demonstrated in previous experimental and computational studies [5], [35]. In *Fo*FaeC, Ser237, located in a position relevant to Arg411 of *Is*MHETase (Table S3), provides more space for the accommodation of longer ligands in the distal pocket of the active site, possibly explaining the ability of *Fo*FaeC WT to also act on BHET and other PET oligomers. Also, in *Fo*FaeC_G122S, the aromatic ring of Phe342 which overlaps with the Gln410 *Is*MHETase amide group, may enhance substrate selectivity towards longer phenolic compounds through potential π-π stacking interactions. Compared to Ser131 of *Is*MHETase, the hydroxyl group of engineered Ser122 in *Fo*FaeC_G122S is shifted to form a hydrogen bond with the alternative conformation of the catalytic Ser201 (Fig. 3B). Conversely, Ser131 in *Is*MHETase points towards the protein surface, where it is exposed to the solvent molecules (Fig. S4). A cross-substitution of hydrophobic residues occurs within the binding pockets of both enzymes (Fig. 3C), with Phe230 and Ile415 in *Fo*FaeC occupying positions equivalent to Leu254 and Phe495 in *Is*MHETase, respectively.

**Figure 3.**
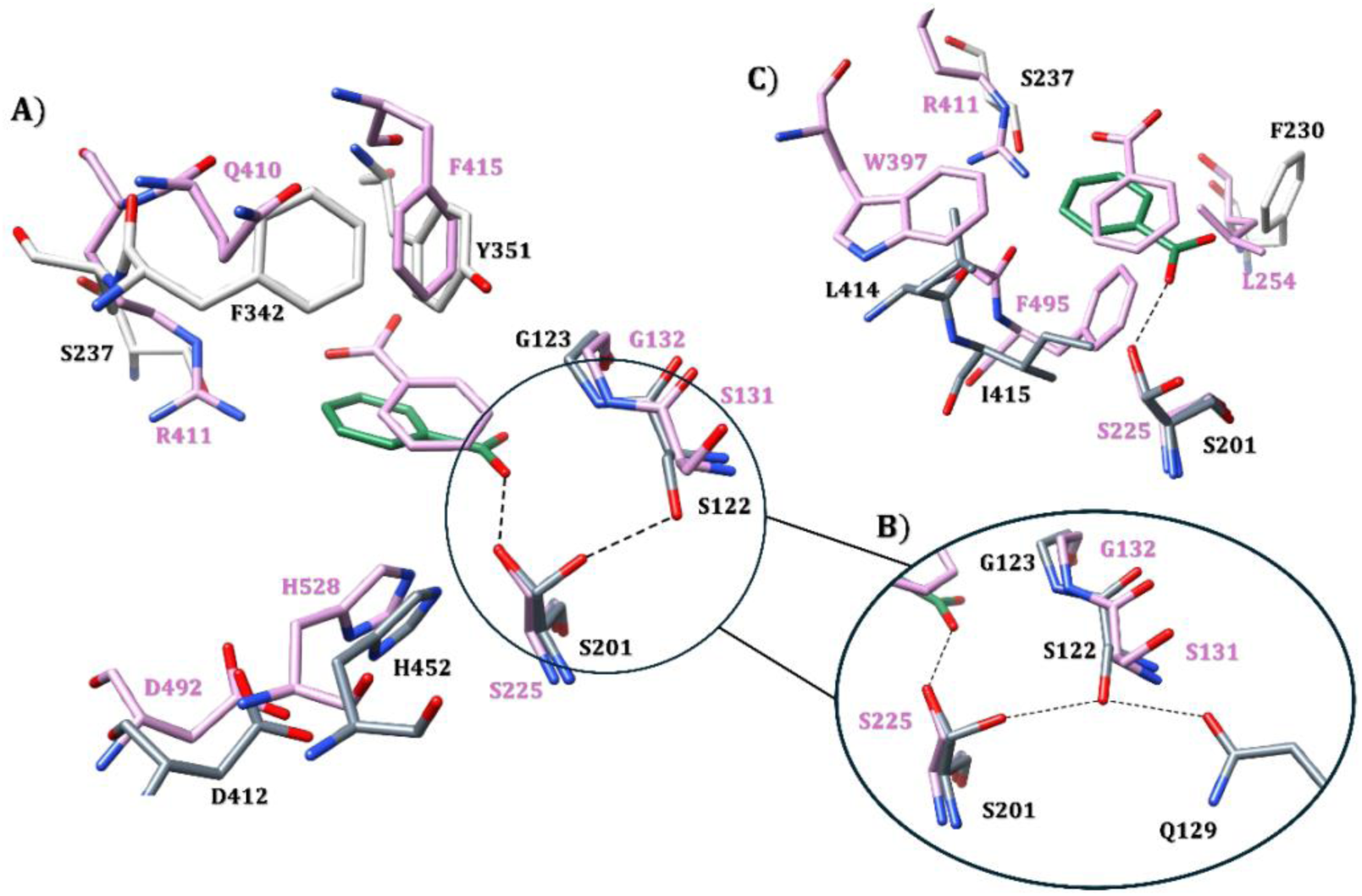
Comparison of FoFaeC_G122S and IsMHETase BA binding site. (A) Global view of the catalytic region of FoFaeC_G122S (light/dark gray) and IsMHETase-6QZ3 (pink), in complex with BA (B) Close-up view of engineered S122 and (C) Hydrophobic cleft of binding pocket of the two enzymes.

### Molecular dynamics and docking simulations

#### MD simulations of apo structures

MD simulations with a total duration of 4 μs (2 systems x 4 trajectories x 500 ns) were used to estimate enzyme behavior before and after introducing mutation G122S. Both WT and MT are quite stable throughout the simulation, with an RMSD value of about 0.15 nm (Fig. S5). Additionally, thermostability analysis using the dimensionless parameter *Lambda* (Λ) indicated that the engineered mutation does not affect the overall thermostability of the enzyme, as the maximum distribution of both WT and MT occurs at the same Λ value (Fig. S6). This suggests that the backbone N-H vectors are temperature independent (within the temperature range probed), in agreement with the biochemical findings reported by *Makryniotis et al.* [9].

The most important alterations apparently induced by G122S mutation are observed in regions 70-75, 96-100, 120-130, 142-179 and 295-306 (Fig. 4 and S7). With the exception of residues 70-75 (loop 3), which belong to the hydrolase domain and seem to become more flexible after the engineered mutation, all other amino acid groups appear to become less flexible. The region 120-130 (loop 6) contains the engineered mutation, and its increased stability is apparently due to the hydrogen bonds introduced by the hydroxyl group of Ser122. Reduced flexibility is also observed at Gly123 of the oxyanion hole of the binding pocket and at the peptide bond Ala125-Gly126, which adopts *trans/cis* conformation in crystallographic MT structure. This region is located next to the catalytic serine, potentially contributing to substrate binding during catalysis. The enhanced stabilization of loop 7 (residues 142-179) is also associated with the *trans* to *cis* rearrangement, mediated by the hydrogen bond between Ala125 and Asn148 observed in the *cis* conformation. Last, loop 10 (residues 295-306) corresponds to a highly flexible loop on the upper part of the lid domain, evidenced from high B-factors relative to the rest of the structure, but also the absence of sufficient electron density in the crystallographic structure of *Fo*FaeC in complex with *p*CA acid (PDB code: 8BHH). The stabilization of this loop is also observed in the case of *Is*MHETase upon substrate binding [35]. Specifically, MD simulations of apo- and substrate bound *Is*MHETase revealed a substrate-induced stabilization of loop 14 (residues 325-351 in *Is*MHETase, 288-307 (loop 10) in *Fo*FaeC) and loop 18 (residues 454-462 in *Is*MHETase). In *Fo*FaeC, the region corresponding to loop 18 of *Is*MHETase harbors an extended N-glycosylation from Asn362.

**Figure 4.**
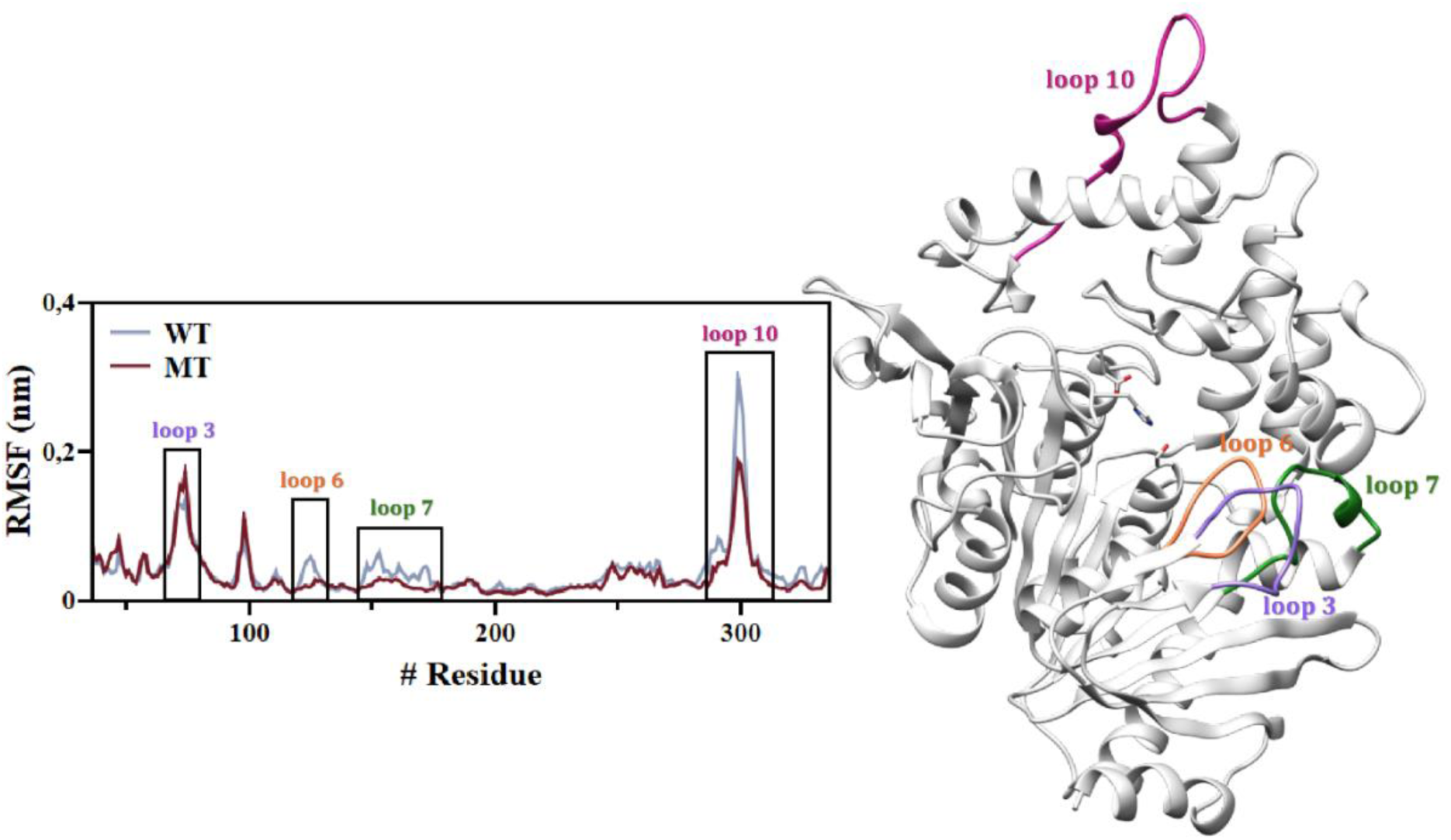
RMSF profiles of WT and MT FoFaeC over 500 ns MD simulation. The WT FoFaeC is shown as a light blue line, while the MT is shown as a dark red line. Loops 3, 6, 7 and 10 are highlighted in purple, orange, green and magenta, respectively, in both the RMSF diagram and the cartoon representation of FoFaeC. The catalytic triad (Ser201 – His452 – Asp412) is depicted as light gray sticks and colored by heteroatoms (oxygen – red, nitrogen – blue). The full RMSF plot including all residues (36-543) is provided in Fig. S8.

#### Docking and MD simulations

Molecular docking of *Fo*FaeC and *Fo*FaeC_G122S with MHET was performed to further examine the role of the engineered residue in the increased activity of the enzyme against MHET. Principal component analysis on the motion of the binding pocket residues followed by k-means clustering resulted in four distinct configurations per enzyme (Fig. S9, S10). The binding free energy of the two complexes selected for MD was calculated to be 5.34 kcal mol^− 1^ and 5.36 kcal mol^−1^, for *Fo*FaeC and *Fo*FaeC_G122S respectively, while the dissociation constant for each complex was 0.10 mM and 0.12 mM at 273 K, respectively. The MHET molecule was similarly positioned in both cases, with its aromatic ring being accommodated in the hydrophobic cavity composed of Ile415, Leu414, Phe342 and 349, as well as Tyr351, and its ethylene glycol moiety pointing towards the solvent exposed side of the protein (Fig. 5A). A similar orientation is observed in the MHETA bound structure of *Is*MHETase (PDB code: 6QGA, Fig. 5B). The docked MHET molecule forms hydrogen bonds with catalytic residues Ser201 and His452, as well as with the residues Gly123, Leu233, Gln234, Ser237, Phe349, Leu414 and Ile415 of the binding pocket. In *Fo*FaeC_G122S, MHET additionally forms hydrogen bonds with residues Thr75, Ser122, Met124, Ala227, Phe342 and Tyr351, while in *Fo*FaeC, with residues Thr202 and Cys543.

**Figure 5.**
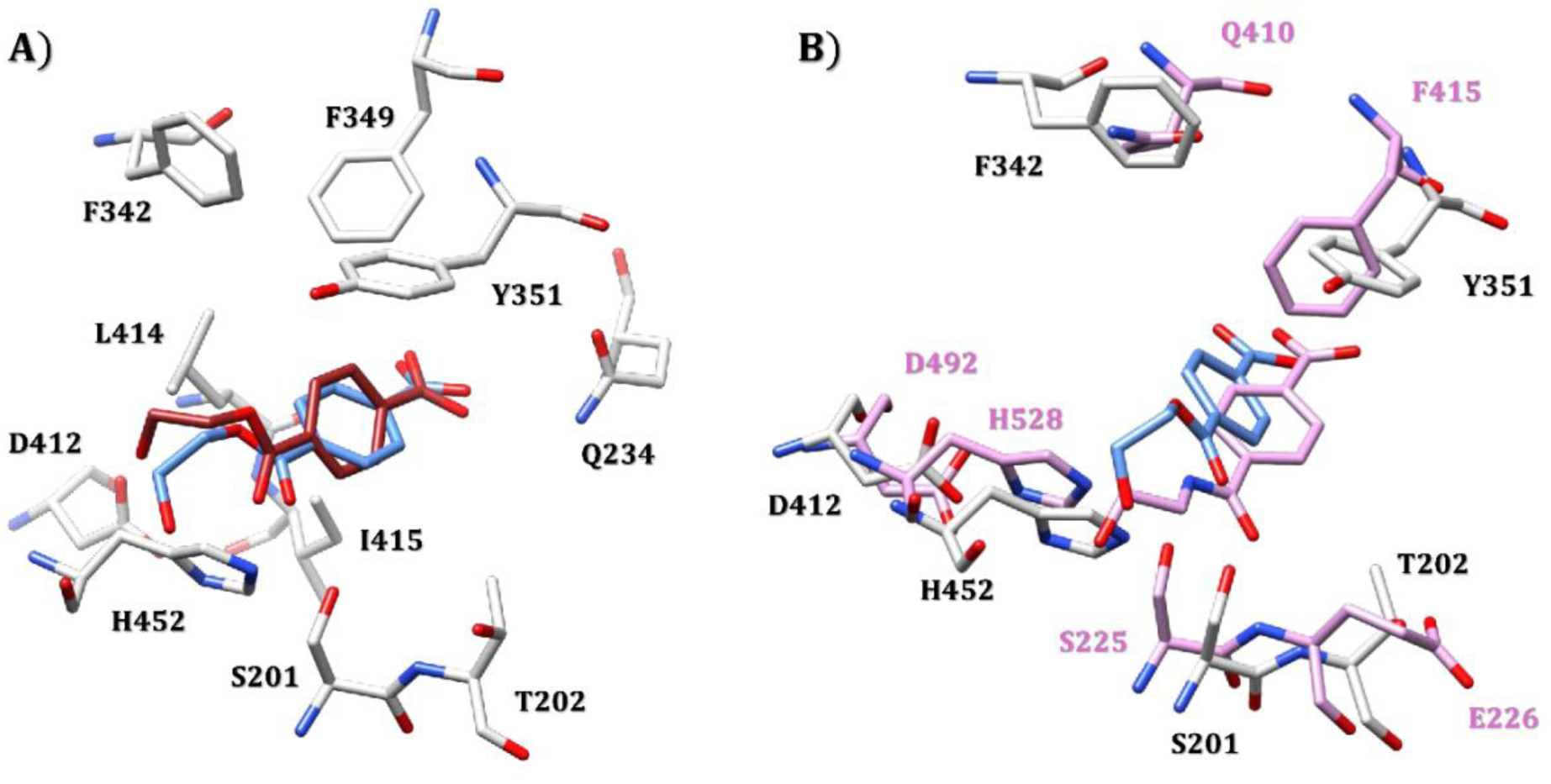
Molecular docking analysis illustrates MHET binding within the active site of *Fo*FaeC. **(A)** Superposition of WT (blue) and MT (red) *Fo*FaeC structures after docking simulations with MHET. **(B)** Superposition of WT *Fo*FaeC (light gray) in complex with MHET (blue) and *Is*MHETase in complex with MHETA (purple). Residues are shown in stick representation and colored by heteroatoms (oxygen – red, nitrogen – blue).

The selected structures (WT and MT in complex with MHET) were equilibrated and subjected to MD dynamics yielding four independent trajectories each, following the same procedure used for the apo structures, increasing the cumulative simulation time for the current study to 8 μs. The *Fo*FaeC and *Fo*FaeC_G122S complexes showed an RMSD value of 0.18 nm and 0.14 nm, respectively, while the RMSD calculated for the carbon atoms of the aromatic ring of MHET moiety remained low at 0.01 nm (Fig. S11). The RMSF analysis of the proteins revealed that loops 6 (the engineered mutation region) and 7 display increased flexibility in the MT complex compared to the apo form, whereas the flexibility of the corresponding loops in the WT remained almost unchanged upon MHET binding (Fig. 6A-6C, S12).

**Figure 6.**
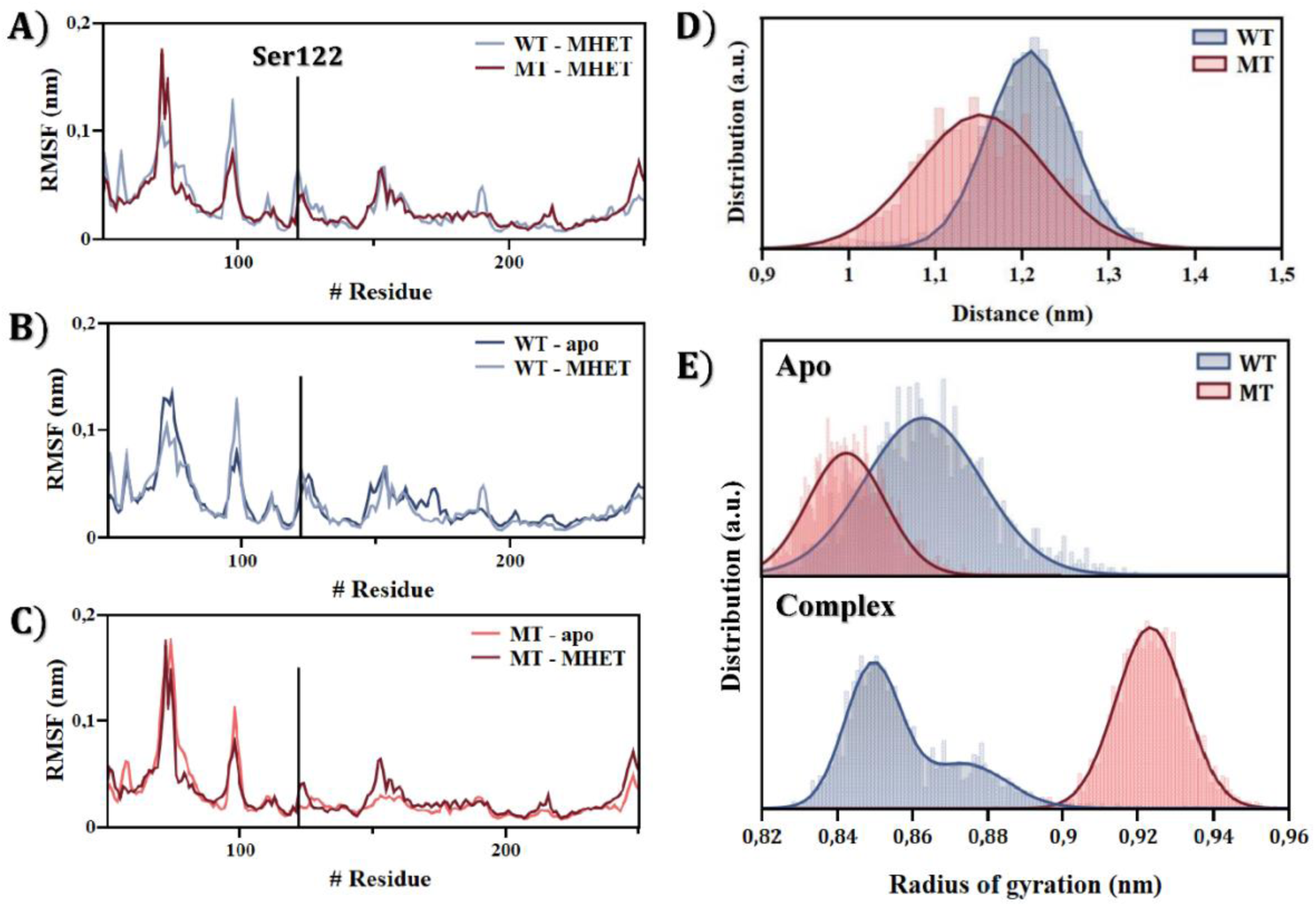
MD analysis. **(A)** RMSF profiles of WT and MT *Fo*FaeC in complex with MHET. **(B)** and **(C)** Comparison of RMSF profiles of apo and MHET-bound forms in WT and MT, respectively. **(D)** Histogram of the distance between Ca atoms of His452 and Phe342, residues of the hydrolase and lid domains, respectively. Distances are calculated for the apo structures. **(E)** Histograms of the radius of gyration of the binding pocket for WT and MT *Fo*FaeC, comparing the apo and MHET-bound states.

During the simulations of the complex structures, the side chain of Phe342 adopts the closed conformation, while Tyr351 exhibited enhanced mobility, forming π-π stacking interactions with the phenolic moiety of the ligand. The observed mobility of Tyr351, as well as the closed conformation of Phe342 in ligand-bound structures, agrees with the crystallographic observation from the *Fo*FaeC_G122S-BA and *Fo*FaeC-*p*CA complex structures. Analysis of the distance between the Ca atoms of His452 (hydrolase domain, catalytic residue) and Phe342 (lid domain) showed that the MT exhibits a broader distance distribution in the unbound state (Fig. 6D), indicating increased relative motion between the hydrolase and lid domains.

In addition to the RMSF analysis, the radius of gyration of the binding pocket was examined for both apo and MHET-bound states for WT and MT *Fo*FaeC. The pocket was defined based on the backbone atoms (N – Ca – C) of the catalytic residues Ser201, His452, Asp412, as well as residues involved in ligand binding, including the engineered Ser122. These residues include Ser122/Gly122, Gly123, Met124, Thr202, Phe230, Leu233, Gln234, Ser237, Phe342, Phe349, Tyr351, Leu414 and Ile415 (Fig. S13). The analysis revealed a noticeably enlarged substrate binding cavity for MT, showing an approximately 2 Å increase in pocket size upon MHET binding (Fig. 6E). This expansion suggests that MT binding pocket can accommodate larger substrates, in agreement with the biochemically observed promiscuity of the enzyme [9].

## Conclusions

The efficient degradation of PET by the plastic degrading bacterium *Ideonella sakaiensis* 201-F6 requires the synergistic effect of PETase and MHETase, highlighting the necessity of the later for efficient plastic degradation to its monomers and subsequent upcycling. One important drawback that has to be tackled before microbe-free enzyme-mediated depolymerization, is substrate inhibition that is observed in MHETase which was alleviated by introducing S131G mutation in *Is*MHETase, which however also reduced reaction efficiency [4]. Here, we present a structural analysis of the “reverse mutant” in *Fo*FaeC, where the corresponding Gly122 was mutated to Ser, resulting in a more efficient and promiscuous enzyme variant.

Despite co-crystallizing *Fo*FaeC_G122S with MHET, clear electron density next to the catalytic serine indicated the presence of BA. The crystal structure of *Is*MHETase in complex with BA has also been previously reported [4], [5], however, it is not clear whether these structures are derived from crystals soaked or co-crystallized with BA. One explanation for the observation of benzoic instead of MHET in *Fo*FaeC_G122S active site could be radiation damage during X-ray data collection [36].

The analysis of the biochemical characteristics of *Fo*FaeC_G122S revealed a more promiscuous enzyme, capable of catalyzing the breakage of a greater variety of natural and synthetic esters [9]. The crystal structure of *Fo*FaeC_G122S in apo state reveals a twisted *trans* conformation in the peptide bond between amino acids 125 and 126 in monomer A, and *cis* conformation in monomer B. Also, interestingly, and in contrast to the WT-apo enzyme, such transition to *cis* peptide is also observed in the *p*CA acid bound structure of *Fo*FaeC WT (PDB code: 8BHH) and the BA bound structure of *Fo*FaeC_G122S. In parallel, MD simulations reveal that the engineered mutation results in a less flexible loop L6, which is the loop that harbors the aforementioned amino acids, as well as loop10, which has been shown to be stabilized upon MHET binding in *Is*MHETase. Gly126 is a conserved amino acid among *Fo*FaeC homologues (Fig. S14). Given the inherent flexibility of glycine and its ability to accommodate backbone conformational changes, its conservation suggests a structural role in modulating the geometry required for substrate binding. The implication of this region to substrate stabilization has been recently also demonstrated in the computational work by *Wang et al* [35]. Additionally, Phe342 is found in two distinct conformations in the determined crystal structures, an open state which is combined with a *trans* conformation of 125-126 bond and a closed one, found in complex structures, and combined with the *cis* conformation observed in the 125-126 peptide bond. It could thus be suggested that the introduced mutation favours a structural rearrangement of the enzyme’s active site, to resemble the ligand-bound state, involving the movement of Phe342 acid chain and *cis-trans* alteration of 125-126 peptide bond and the stabilization of loops 6 and 10. The implication of this *cis-trans* variation in ligand binding has been shown in previous studies [37], and suggested to be related to enzyme evolution for new substrates and functions. The role of Gln410 of *Is*MHETase, located at a position corresponding to Phe342 has been shown in the work by *Knott et al* [4], who suggested a concerted movement of Phe415 that adopts an open and close conformation, and Gln410 upon substrate binding. Similarly, Tyr351 of *Fo*FaeC, which occupies a position analogous to Phe415 of *Is*MHETase, has been reported in previous X-ray crystallographic studies to exhibit high mobility and to contribute to substrate binding through the formation of π-π stacking interactions with the aromatic rings of the substrates [8].

In summary, the introduced mutation yields a variant with broadened substrate scope, driven by the formation of novel intramolecular interactions, loop stabilization, and expansion of the binding cavity. This study advances our understanding of the structural determinants underlying increased enzyme promiscuity in an MHET-active esterase. The insights obtained provide a foundation for future enzyme-engineering strategies aimed at developing robust biocatalysts for a wide range of biotechnological applications, including plastic biodegradation. Moreover, in an era of rapidly advancing computational structure prediction, this work underscores the value of integrating experimental and computational approaches, particularly for validating the structural and functional consequences of point mutations.

## Data availability

The atomic coordinates of the three-dimensional structures presented in this work have been deposited in the Protein Data Bank (PDB) with PDB ID: 9I50 and 9HUN.

## Abbreviations

a.u: asymmetric unit
BA: benzoic acid
BHET: bis(2-hydroxyethyl) terephthalate
CAZy: Carbohydrate-Active Enzymes database
EG: ethylene glycol
FA: ferulic acid
FAE: ferulic acid esterase
*Fo*FaeC: *Fusarium oxysporum* FAE
*Fo*FaeC_G122S: *Fo*FaeC variant G122S
*Is*MHETase: *Ideonella sakaiensis* MHETase
MCA: methyl caffeate
MD: molecular dynamics
MFA: methyl ferulate
MHET: mono(2-hydroxyethyl) terephthalate
M(HET)_2_: a soluble methylated PET dimer
MHETase: mono(2-hydroxyethyl) terephthalate esterase
M*p*CA: methyl *p*-coumarate
MT: mutant
*p*CA: *p*-coumaric acid
PCA: Principal Component Analysis
PET: polyethylene terephthalate
PETase: polyethylene terephthalate esterase
RMSD: root mean square deviation
RMSF: root mean square fluctuation
TPA: terephthalic acid
WT: wild-type

## Acknowledgements

This research was funded by the Hellenic Foundation for Research and Innovation (H.F.R.I.) under the “Basic Research Funding (Horizontal support for all Sciences), National Recovery and Resilience Plan (Greece 2.0) - Sub-action II Funding Projects in Leading-Edge Sectors” (EnZyReMix, project number: 15024). The synchrotron data was collected at beamline P13 operated by EMBL Hamburg at the PETRA III storage ring (DESY, Hamburg, Germany). We would like to thank Dr. Isabel Bento for the assistance in using the beamline. This work was also supported by computational time granted from the National Infrastructures for Research and Technology S.A. (GRNET S.A.) in the National HPC facility - ARIS - under project ID “MultiEng”.

## Author contributions

MD and ET conceptualization. MD and VD methodology. PK, TIS, MD and VD data curation and formal analysis. PK and KM performed the experiments. PK and MD writing original draft. PK, KM, ET, VD, MD writing – review and editing. MD, ET and VD funding acquisition. MD and VD supervison.

