## Supplementary_data for "Structural investigation of an engineered feruloyl esterase with improved MHET degrading properties"

### Supplementary information

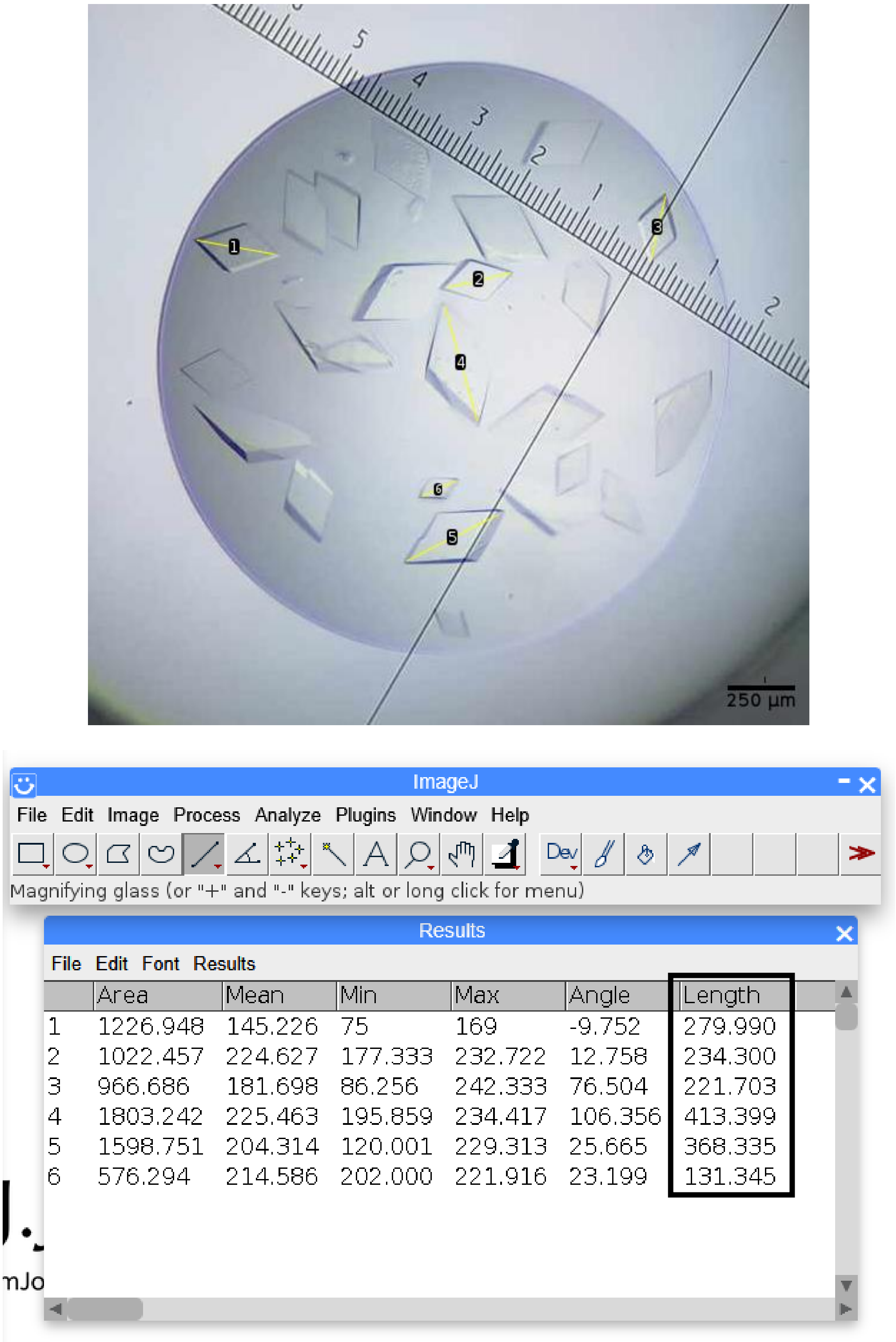


| | Area | Mean | Min | Max | Angle | Length |
|---|---|---|---|---|---|---|
| 1 | 1226.948 | 145.226 | 75 | 169 | -9.752 | 279.990 |
| 2 | 1022.457 | 224.627 | 177.333 | 232.722 | 12.758 | 234.300 |
| 3 | 966.686 | 181.698 | 86.256 | 242.333 | 76.504 | 221.703 |
| 4 | 1803.242 | 225.463 | 195.859 | 234.417 | 106.356 | 413.399 |
| 5 | 1598.751 | 204.314 | 120.001 | 229.313 | 25.665 | 368.335 |
| 6 | 576.294 | 214.586 | 202.000 | 221.916 | 23.199 | 131.345 |

***Figure S1.*** *FoFaeC_G122S crystals grown in the presence of 0.1 M Tris-HCl pH 8.5 and 35% (v/v) PEG400. The size of the crystals were measured using ImageJ online program ( https://imagej.net/ij/ ).*

***Table S1.*** *Data collection and processing statistics.*

*Values for the outer shell are given in parentheses.*

| PDB code | 9I50 | 9HUN |
|---|---|---|
| Diffraction source | P13, PETRA III | P13, PETRA III |
| Wavelength (Å) | 0.9763 | 0.9763 |
| Temperature (K) | 100 | 100 |
| Detector | DECTRIS EIGER X 16M | DECTRIS EIGER X 16M |
| Crystal-detector distance (mm) | 296.31 | 297.44 |
| Rotation range per image (°) | 0.01 | 0.01 |
| Total rotation range (°) | 360 | 360 |
| Exposure time per image (s) | 0.1 | 0.1 |
| Space group | $P2_1$ | $P2_1$ |
| *a*, *b*, *c* (Å) | *68.56, 89.26, 116.48* | *68.00, 89.93, 115.15* |
| α, β, γ (°) | 90, 101.72, 90 | 90, 103.77, 90 |
| Resolution range (Å) | 114.05 – 1.90 (1.93 – 1.90) | 89.93 – 1.71 (1.74 – 1.71) |
| Total No. of reflections | 751338 (38439) | 491252 (24257) |
| No. of unique reflections | 107339 (5299) | 143024 (7131) |
| Completeness (%) | 99.2 (99.6) | 98.6 (99.2) |
| Half-set correlation CC(1/2) | 0.999 (0.739) | 0.995 (0.812) |
| Redundancy | 7.0 (7.3) | 3.4 (3.4) |
| $\langle I/\sigma(I)\rangle$ | 13.0 (1.6) | 7.5 (1.5) |
| $R_{merge}$ | 0.077 (1.241) | 0.065 (0.537) |
| $R_{\text{p.i.m.}}$ | 0.048 (0.752) | 0.062 (0.493) |
| Overall *B* factor from Wilson plot (Å$^2$) | 31.89 | 28.71 |

***Table S2.*** *Structure solution and refinement.*

*Values for the outer shell are given in parentheses.*

| PDB code | 9I50 | 9HUN |
|---|---|---|
| Final $R_{cryst}$ | 0.162 | 0.161 |
| Final $R_{free}$ | 0.197 | 0.193 |
| σ cutoff | none | none |
| No. of reflections, working set | 107314 | 142930 |
| No. reflections, test set | 5438 | 7141 |
| No. of protein molecules in asymmetric unit | 2 | 2 |
| No. of non-H atoms | | |
| Protein | 7981 | 8057 |
| Ion | 2 | 2 |
| Ligand | 612 | 671 |
| Water | 606 | 814 |
| Total | 9201 | 9544 |
| R.m.s. deviations | | |
| Bonds (Å) | 0.0071 | 0.0131 |
| Angles (°) | 1.582 | 1.990 |
| Average *B* factors (Å$^2$) | | |
| Protein | 43.90 | 44.21 |
| Ion | 38.43 | 41.22 |
| Ligand | 59.36 | 58.83 |
| Water | 48.72 | 50.93 |
| Ramachandran plot | | |
| Most favoured (%) | 97.33 | 97.92 |
| Outliers (%) | 0.40 | 0.49 |

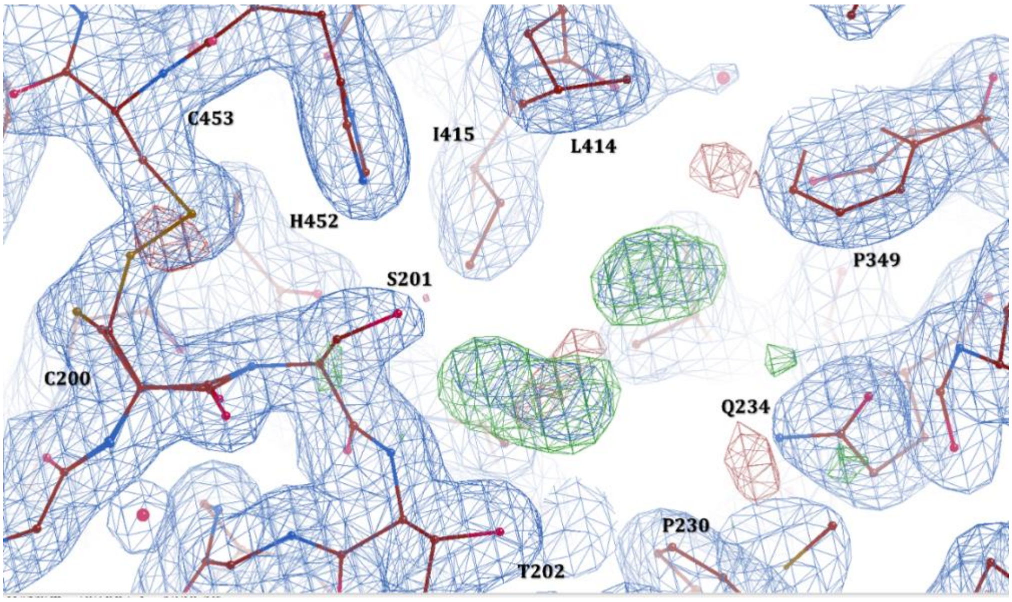


***Figure S2.*** *Representation of the crystal structure of FoFaeC_G122S, chain B (PDB ID: 9I50) in COOT. Amino acids are shown in stick models colored by heteroatoms. The* $2F_o - F_c$ *electron density map is contoured at 1σ (blue mesh), while the* $F_o - F_c$ *difference map is contoured at 3σ (green mesh).*

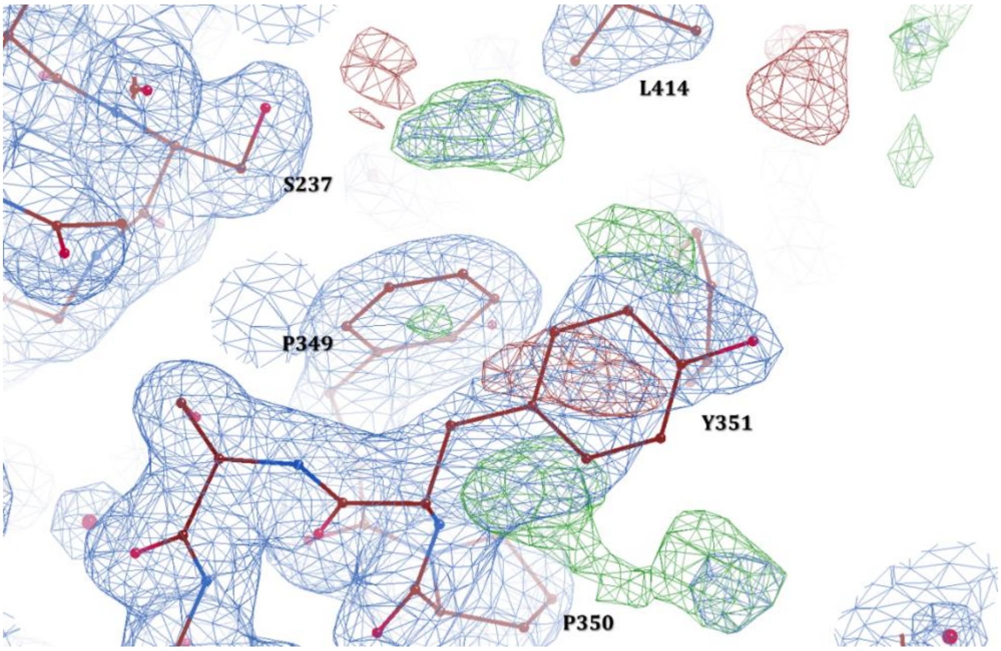


***Figure S3.*** *Representation of the crystal structure of FoFaeC_G122S, chain B (PDB ID: 9I50) in COOT. Close-up view of Tyr351 highlighting its mobility due to the non-satisfied electron density map around it. Amino acids are shown in stick models colored by heteroatoms. The* $2F_o - F_c$ *electron density map is contoured at 1σ (blue mesh), while the* $F_o - F_c$ *difference map is contoured at 3σ (green mesh).*

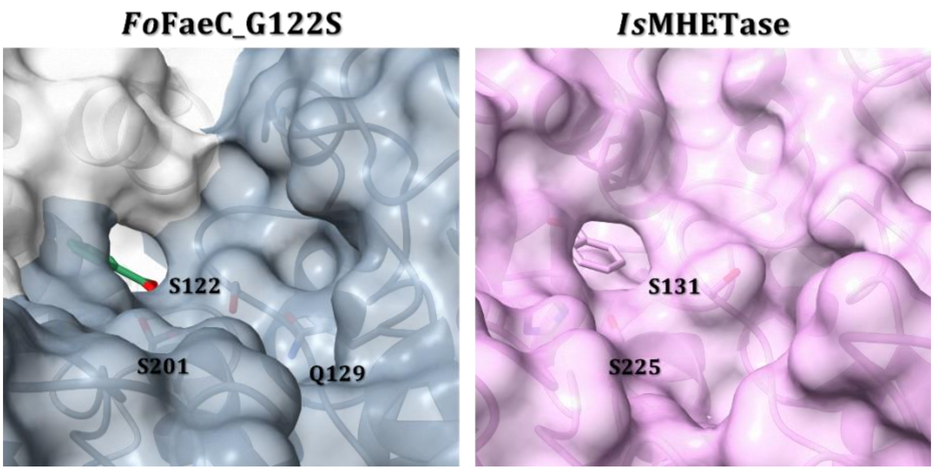


***Figure S4.*** *Surface representation of the cavity around hydroxyl group of Ser131 in IsMHETase (PDB code 9QZ3) and engineered Ser122 in FoFaeC_G122S-BA. IsMHETase is shown in pink color, while FoFaeC_G122S is in gray colors. Ser131 of IsMHETase interacts with water molecules, while Ser122 in FoFaeC_G122S-BA forms hydrogen bonds with Gln129 and catalytic Ser201.*

***Table S3.*** *Residues implicated in catalysis and substrate binding, in FoFaeC and IsMHETase. Those belonging to the lid domain are shown in red, while those from the catalytic domain in blue.*

| | ***Fo*FaeC** | ***Is*MHETase** |
|---|---|---|
| **Enzyme catalytic** | S201 | S225 |
| | D412 | D492 |
| | H452 | H528 |
| **Oxyanion hole** | G123 | G132 |
| | T202 | E226 |
| **Gate Residue** | Y351 | F415 |
| **BHET binding residues** | F230 | L254 |
| | L414 | W397 |
| | S237 | R411 |
| | Y351 | S416 |
| | I415 | F495 |
| | G123 | G132 |
| | L414 | A494 |
| **BA** | F342 | Q410 |
| **Mutation** | S122 | S131 |

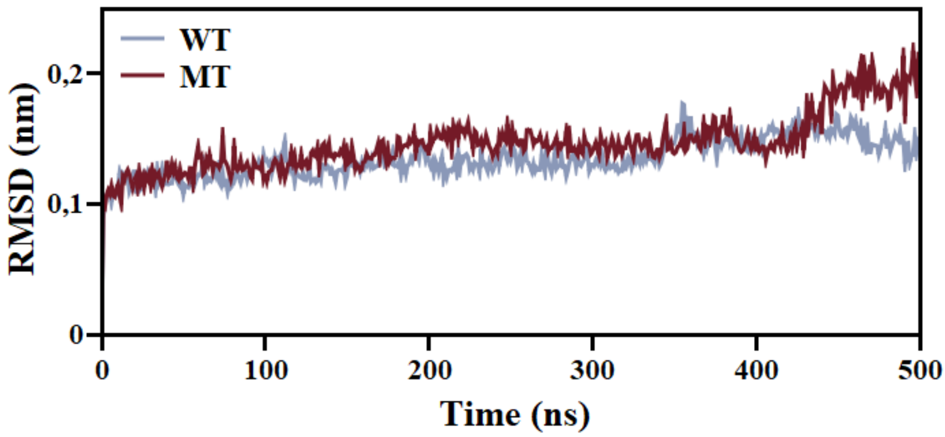


***Figure S5.*** *Root Mean Square Deviation (RMSD) profiles of FoFaeC over 500 ns of molecular dynamics simulation. The wild-type (WT) FoFaeC is shown in blue, while the mutant (MT) FoFaeC_G122S is in red.*

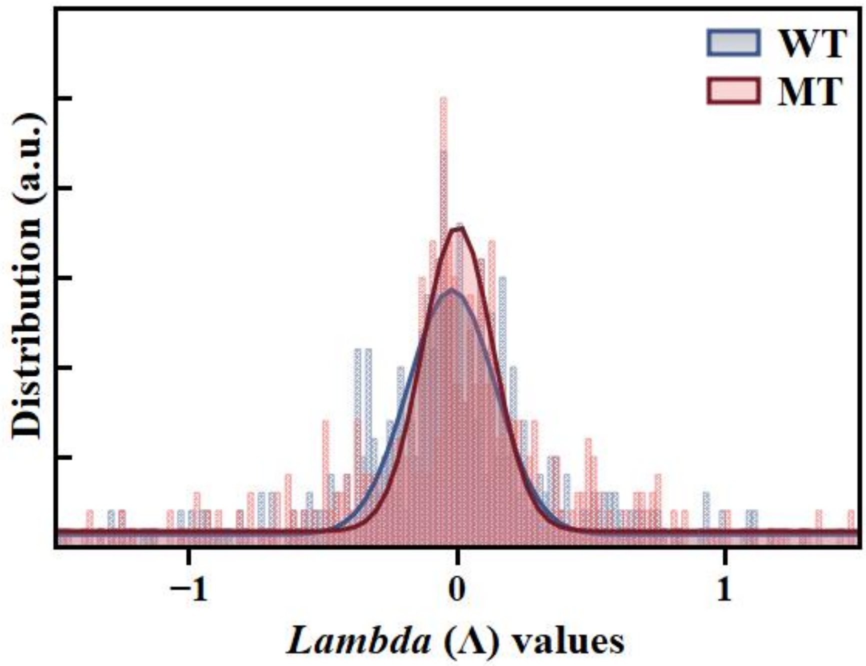


***Figure S6.*** *Histogram of Λ values for the wild-type and mutant G122S for the whole enzymes. A solid line guide (Gaussian fitting) is provided for each histogram to determine the Λ value of the maximum distribution of each enzyme. Lower Λ values indicate higher protein thermostability.*

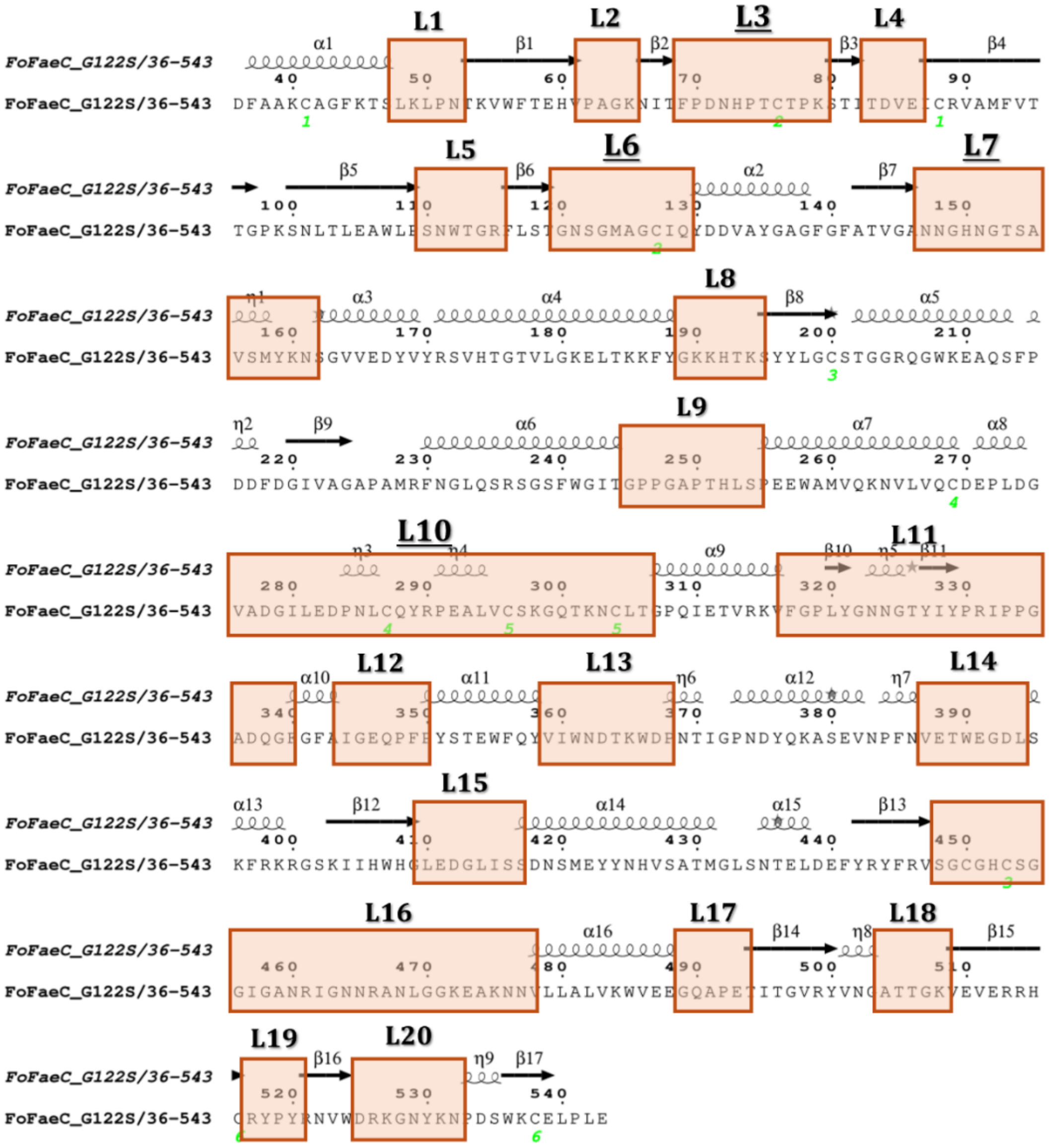

***Figure S7.*** *Secondary structure elements correspond to FoFaeC_G122S, highlighting loops (L) regions (L1-20). It also shows the α-helices and β-strands. The figure was prepared using ESPRIPT (https://espript.ibcp.fr/ESPript/ESPript/).*

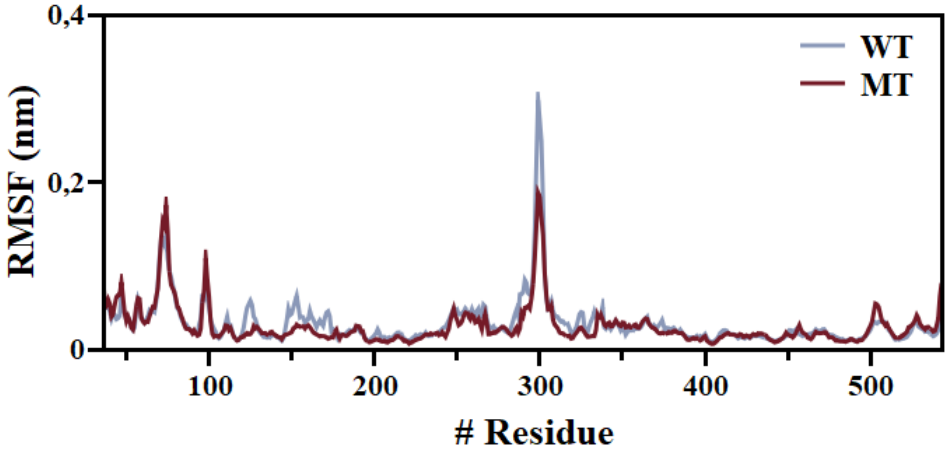


***Figure S8.*** *Root Mean Square Fluctuation (RMSF) profiles of FoFaeC over 500 ns molecular dynamics simulation. The wild-type (WT) FoFaeC is shown in blue, while the mutant (MT) FoFaeC_G122S is in dark red.*

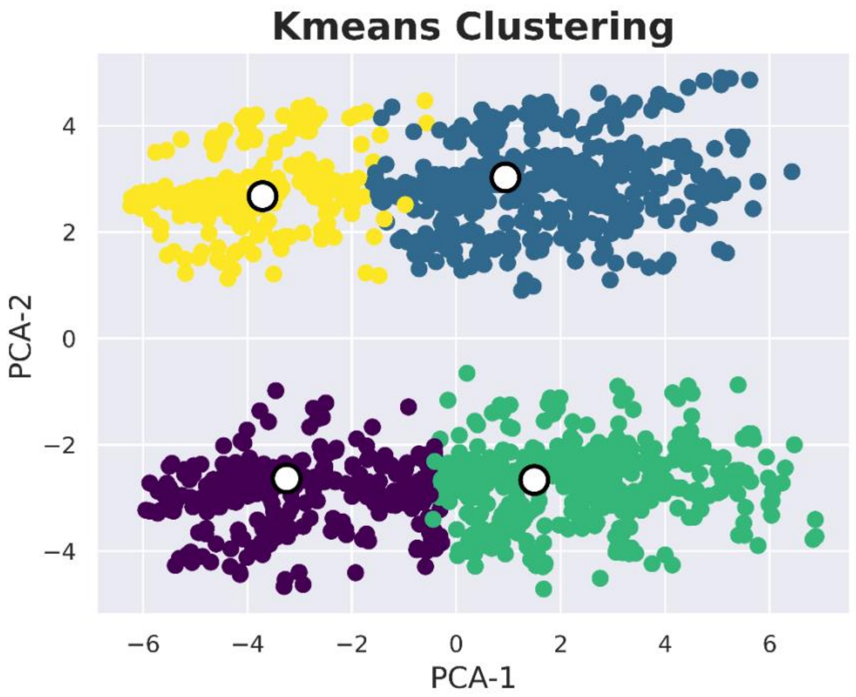


***Figure S9.*** *Total K-means clustering of the four independent WT trajectories over 500 ns MD simulation.*

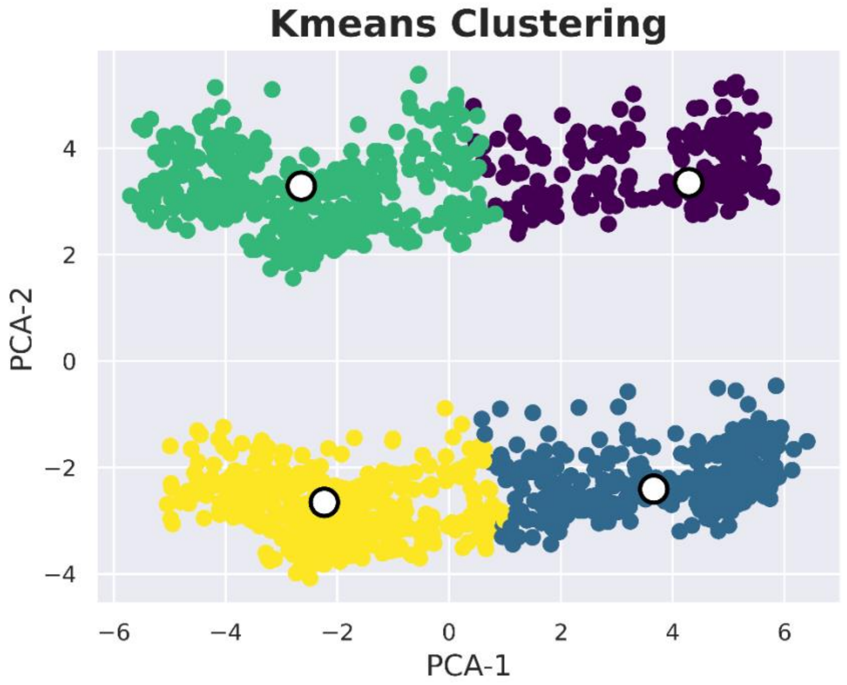


***Figure S10.*** *Total K-means clustering of the four independent MT trajectories over 500 ns MD simulation.*

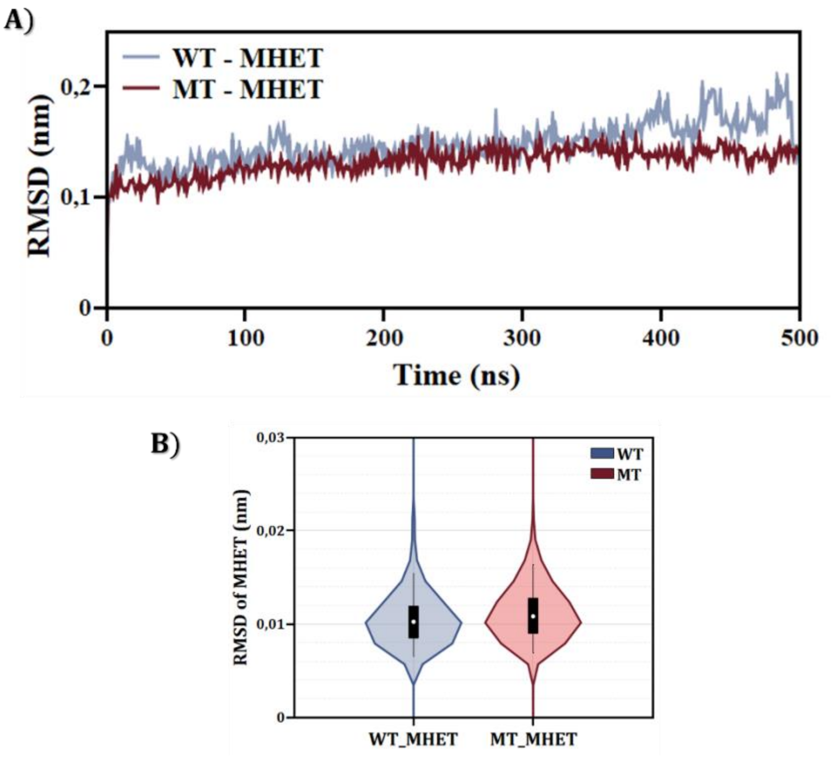


***Figure S11.*** *RMSD profiles of FoFaeC (A) and violin plots of the RMSD for the aromatic ring of MHET (B) over 500 ns of molecular dynamics simulation. The wild-type (WT) FoFaeC is shown in blue, while the mutant (MT) FoFaeC_G122S is in red.*

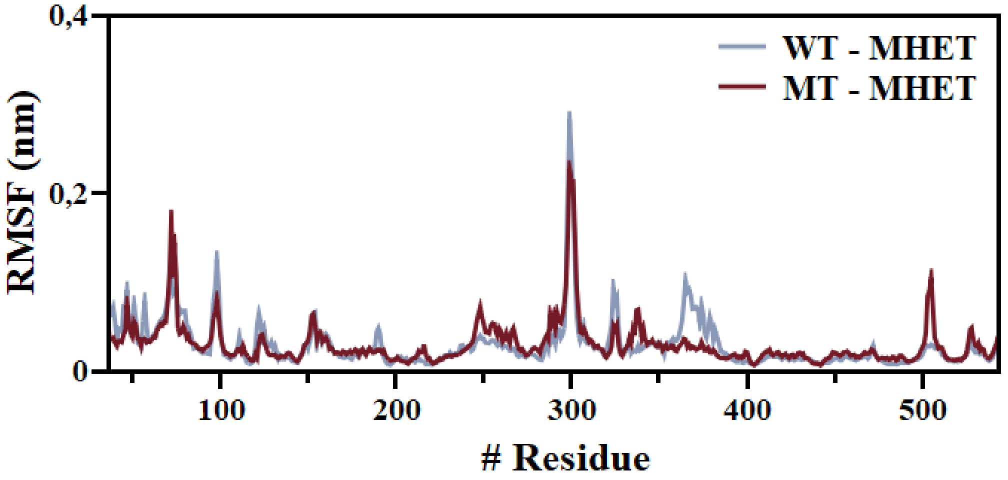


***Figure S12.*** *Root Mean Square Fluctuation (RMSF) profiles of FoFaeC in complex with MHET over 500 ns molecular dynamics simulation. The wild-type (WT) FoFaeC is shown in blue, while the mutant (MT) FoFaeC_G122S is in dark red.*

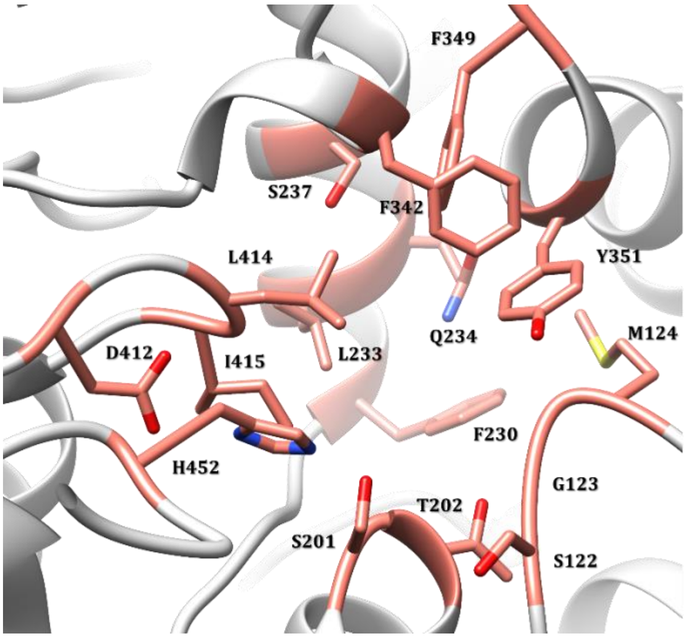


***Figure S13.*** *Residues forming the binding pocket in MT and used for the $R_g$ calculation are shown in pink stick representation.*

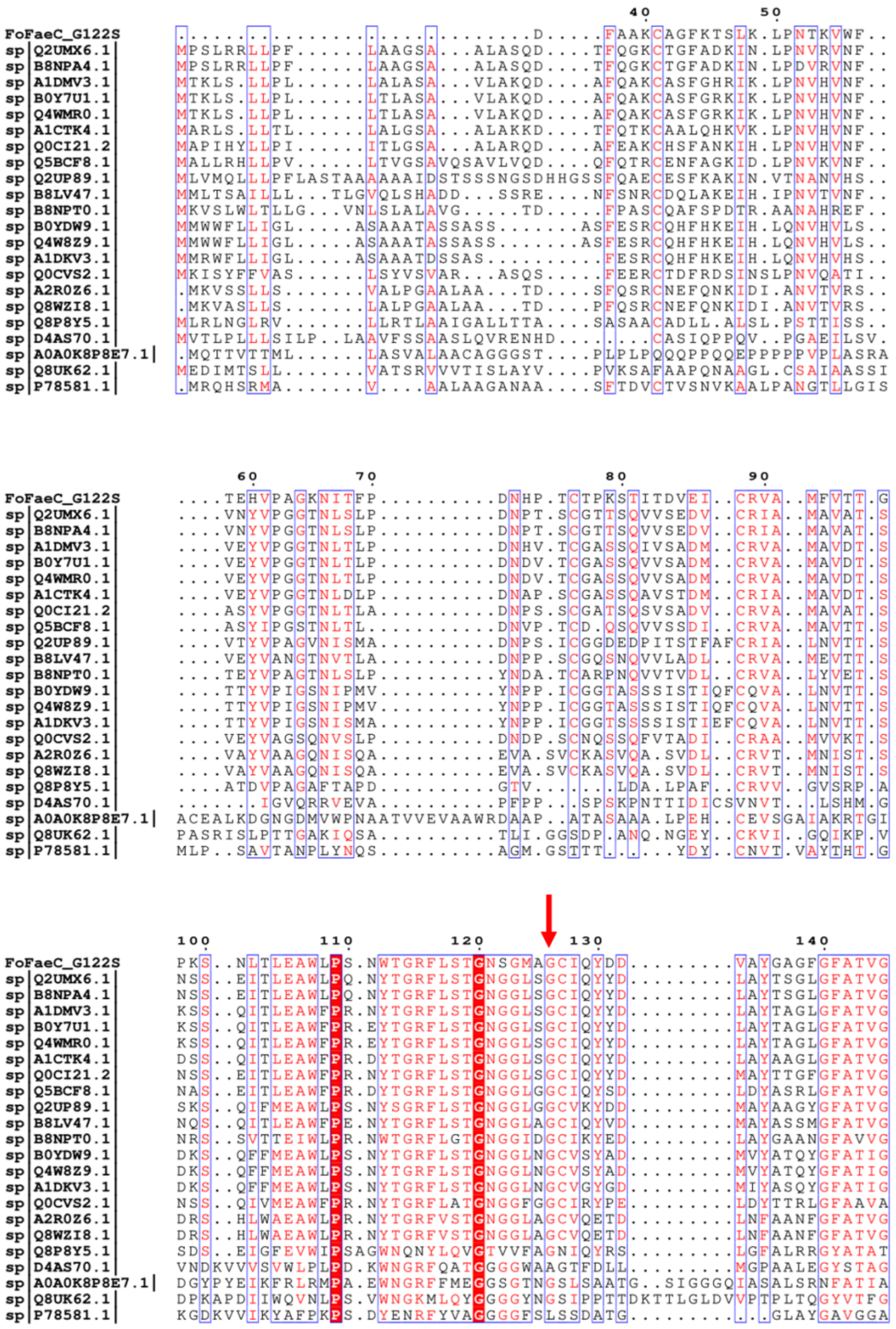

***Figure S14.*** *A BLAST search against UniProtKB/Swiss-Prot database was performed using the FoFaeC_G122S sequence as input. To identify evolutionary conserved regions, the BLAST output FAE sequences were aligned with FoFaeC_G122S, revealing that Gly126 is highly conserved among the homologous proteins (red arrow). The figure was prepared using ESPRIPT (*[*https://espript.ibcp.fr/ESPript/ESPript/*](https://espript.ibcp.fr/ESPript/ESPript/)*).*
